# Polygenic prediction of type 2 diabetes in continental Africa

**DOI:** 10.1101/2021.02.11.430719

**Authors:** Tinashe Chikowore, Kenneth Ekoru, Marijana Vujkovic, Dipender Gill, Fraser Pirie, Elizabeth Young, Manjinder S Sandhu, Mark McCarthy, Charles Rotimi, Adebowale Adeyemo, Ayesha Motala, Segun Fatumo

## Abstract

**Objective:** Polygenic prediction of type 2 diabetes in continental Africans is adversely affected by the limited number of genome-wide association studies (GWAS) of type 2 diabetes from Africa, and the poor transferability of European derived polygenic risk scores (PRS) in diverse ethnicities. We set out to evaluate if African American or multi-ethnic derived PRSs would improve polygenic prediction in continental Africans.

**Research Design and Methods:** Using the PRSice software, ethnic-specific PRSs were computed with weights from the type 2 diabetes GWAS of the Million Veteran Program (MVP) study. The South African Zulu study (1602 cases and 976 controls) was used as the target data set. Replication and assessment of the best predictive PRS association with age at diagnosis was done in the Africa America Diabetes Mellitus (AADM) study (1031 cases and 738 controls).

**Results:** The African American derived PRS was more predictive of type 2 diabetes compared to the European and multi-ethnic derived scores. Notably, participants in the 10^th^ decile of this PRS had a 3.19-fold greater risk (OR 3.19; 95%CI (1.94-5.29), p = 5.33 x10^-6^) of developing diabetes and were diagnosed 2.6 years earlier compared to those in the first decile.

**Conclusions:** African American derived PRS enhances polygenic prediction of type 2 diabetes in continental Africans. Improved representation of non-Europeans populations (including Africans) in GWAS, promises to provide better tools for precision medicine interventions in type 2 diabetes.

## Introduction

The global prevalence of diabetes mellitus in 2019 was estimated to be 463 million individuals(1), of which 19.4 million are from Africa and 90% of them had type 2 diabetes. African countries are adversely affected by limited resources to manage this burden. Nonetheless, by 2045 it is projected that Africa will experience the largest increase of diabetes prevalence in the world of 143% (1; 2). In addition, the highest proportion of undiagnosed (59.7%) people living with diabetes in the world reside in Africa(1). Therefore, urgent strategies and resources for improving screening and early identification interventions are required to help curb this pandemic in Africa.

Type 2 diabetes is a multifactorial disease, that is hypothesised to be increasing in prevalence due to the interaction of genetic and environmental factors(3). Although the genetic factors are stable overtime, the surge in diabetes prevalence over the past decades is thought to be caused by urbanization, and adoption of westernized lifestyles characterised by consumption of energy dense foods and physical inactivity(3; 4). However, diabetes has been noted to be preventable and its onset delayed for 15 years by diet and exercise interventions in the Diabetes Prevention Program(5). Since diet and exercise strategies are readily accessible and relatively low-cost, coupling these lifestyle interventions with approaches that identify people more susceptible to develop diabetes earlier, might effectively lower the diabetes burden. The use of polygenic risk scores for early identification of people that are more genetically susceptible to develop type 2 diabetes is such an approach(6). Recent studies conducted in Europeans have indicated that individuals in the 10^th^ decile have 5.21-fold higher risk (OR=5.21; 95% CI 4.94–5.49) of developing diabetes compared to those in the first decile(7). However, evidence exists of the poor transferability European derived polygenic scores in diverse populations. For example, Martin et al. 2019 reported that European PRSs had a 4.9-fold reduced predictive in Africans Americans across 17 traits. There is a now a concern that African ancestry and other similarly under-studied population groups may not benefit from the clinical translation efforts of these risk polygenic scores and thereby further exacerbate existing health disparities (8; 9).

Large multi-ethnic cohorts such as the Million Veteran Program improve the representation of African Americans in GWAS and offer a promise of improved polygenic prediction in this group (10). However, the representation of continental African in GWAS is still very low, both in the number of studies and the total number of study participants. For example, Type 2 diabetes GWAS studies with over a million Europeans participants are being reported while the sample sizes of continental Africans remain under 10,000 (7; 11). Therefore, continental Africans face a much worse threat than African Americans of under-representation in precision medicine efforts for type 2 diabetes(9). It has been reported that multi-ethnic PRS (compared to European only PRS) might enhance prediction in diverse populations(12; 13). However, the predictive ability of the multi-ethnic derived PRS and that of African Americans (who have ~80% African admixture) is yet to be evaluated in continental Africans (12; 13). We set up this study to evaluate the predictivity of European, African American and multi-ethnic derived polygenic risk scores for type 2 diabetes in continental Africans.

## Methods

### Study participants

Participants were from the Durban Case Control (DCC) study (1602 cases) and 976 controls from the Durban Diabetes Study (DDS) and collectively there are regarded as the South African Zulu study (14). These individuals were above 18 years, not pregnant, and from urban black African communities in Durban, South Africa(14). The WHO criteria was used to define type 2 diabetes status. The replication study participants were from the AADM study which has been described in detail elsewhere(15–17). The 1031 cases and 738 controls from this study were enrolled at university medical centers in Nigeria, Ghana and Kenya(17). In this study, diabetes was defined based on oral glucose tolerance test, fasting glucose or exhibiting symptoms of diabetes(17). Written informed consent was completed by the study participants. The respective studies were approved by relevant ethics committees under the following references DCC (BF078/08), DDS (BF030/12) and AADM (14/WM/1061).

### Genotyping and Imputation

Participants in the South African Zulu study were genotyped using the Illumina Multi-Ethnic Genotyping Array (Illumina, Illumina Way, San Diego, CA, USA). The Affymetrix Axiom PANAFR SNP array or Illumina Multi-Ethnic Genotyping Array was used to genotype participants in the AADM study. Detailed quality control and imputation for these studies has been described elsewhere(11; 18). A minimum MAF threshold of 0.5% and imputation information score > 0.4 was applied(11).

### Statistical Analysis

PRSice 2 software was used to implement the clumping and threshold approach for developing PRS. Clumping distance of 500kb and r2 of 0.5 were parameters used for computing PRS. GWAS summary statistics from the Million Veteran Program (MVP) study(7) were used as the base (discovery) while genotype data from the South African Zulu study and AADM was used as the target data and replication datasets respectively as illustrated in Table 1. The receiver operating curves were computed in R using the pROC package to evaluate the discriminative ability of the PRS from the selected ethinicities. The best predictive PRS was selected based on the Nagelkerke R2 (Supplementary Table 1) and the area under the curve (AUC) see Table 1. Logistic regression models were used to evaluate the predictive capacity of the PRS as shown in Figure 1A and C. These models were corrected for age, sex, body mass index (BMI) and residual population structure using principal components. Finally, a linear regression model was used to evaluate whether age of diagnosis in patients with diabetes (n=1031) is affected by PRS in AADM study, of which the results are shown in Figure 1B.

**Figure 1.**
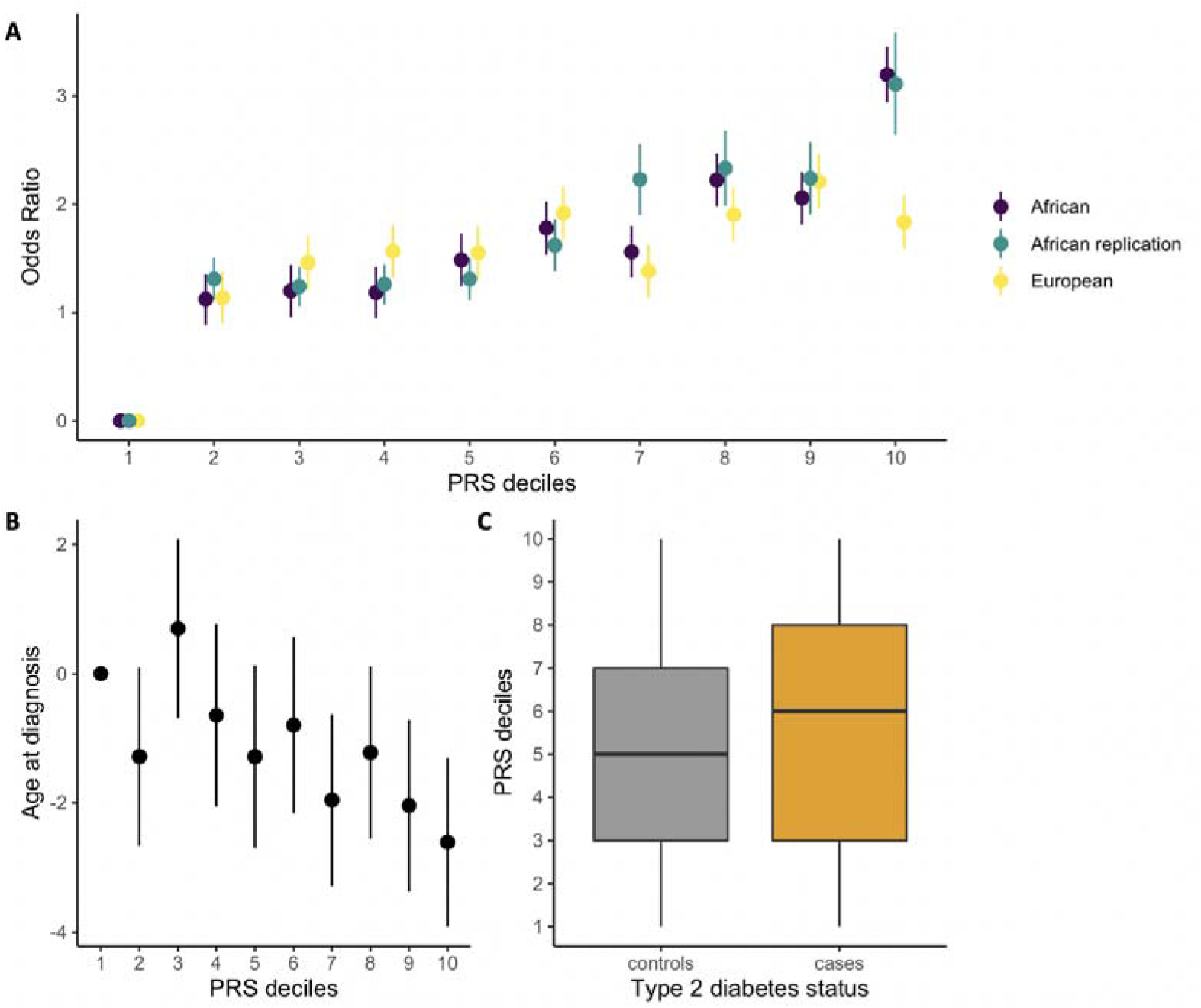
**A.** Shape plot for the difference in odds ratio for type 2 diabetes (adjusted for age, sex, BMI and five principal components) in reference to the 1^st^ decile for the African (African American), European derived PRS in the South African Zulu study and the replication of the African American derived PRS in the AADM study. **B.** Shape plot for the difference of age at diagnosis for type 2 diabetes in the AADM study for the African American derived PRS. **C.** Box plots showing the distribution of the Africa American derived PRS in type 2 diabetes case and controls of the South African Zulu study.

**Table 1.** Comparisons of the predictive ability of ethnically derived PRS on type 2 diabetes in continental Africans.

|  | *Ancestry* |  |  |  |
| --- | --- | --- | --- | --- |
|  | Multiethnic | African American | European | African American Replication |
| ***DiscoveryDataset*** |  |  |  |  |
| Cases | 228,499 | 24,646 | 148,726 | 24,646 |
| Controls | 1,178,783 | 31,446 | 965,732 | 31,446 |
| ***Target Data Set*** |  |  |  |  |
| Cases | 1,602 | 1,602 | 1,602 | 1,031 |
| Controls | 981 | 981 | 981 | 738 |
| ***PRS parameters*** |  |  |  |  |
| P-value threshold | $3 \times 10^{-4}$ | $5 \times 10^{-8}$ | 0.406 | $5 \times 10^{-8}$ |
| Number of SNPs | 41,815 | 65 | 1,441,315 | 60 |
| #AUC % | 56.5 | 58.6 | 56.2 | 58.8 |
| *OR(95%CI) | 1.29 (1.16-1.43) | 1.58 (1.36-1.84) | 1.01 (1.00-1.01) | 1.56 (1.47-1.66) |
| *P-value | $3.52 \times 10^{-6}$ | $4.80 \times 10^{-9}$ | $9.54 \times 10^{-6}$ | $4.81 \times 10^{-21}$ |

\*models adjusted for ancestry indicated by 5 principal components, age, sex and BMI; OR = odds ratio; CI = confidence interval. #AUC = area under the curve; for the PRS adjusted for ancestry indicated by five principal components
\*models adjusted for ancestry indicated by 5 principal components, age, sex and BMI; OR = odds ratio; CI = confidence interval.

## Results

The African American derived PRS had the best predictive capability for type 2 diabetes in continental Africans compared to the European, and the multi-ethnic PRS (Table 1). Comparable odds ratios were noted for this PRS in the South African Zulu study (OR 1.58; p = 4.8 x 10^-9^) and AADM study (OR 1.56, p = 4.81 x 10^-21^) independent of the fact that participants originate from different geographical regions of Africa.

The participants in the 10^th^ decile of the African American derived PRS had a more than 3-fold higher risk for developing type 2 diabetes compared to those in the first decile in both the South African Zulu study (OR = 3.19; p = 5.33 x 10^-6^) and in the AADM study (OR =3.11;p = 7.59 x 10^-14^). Contrariwise, the European derived PRS had a lower predictive ability in continental Africans. The participants in the 10^th^ decile of this PRS had 1.83-fold (OR = 1.83; p = 0.015) higher risk of type 2 diabetes compared to those in the first decile (Figure 1A). On average, participants in the 10^th^ decile of the African American PRS in the AADM study were diagnosed with type 2 diabetes 2.6 years earlier (Beta = −2.61; p = 0.046) than participants in the first decile (Figure 1B).

The median of the African American PRS distribution in the cases was 60% and 50% controls indicating that more cases were in the upper deciles of the score compared to controls in the South African Zulu study as shown in Figure 1C. The model with the conventional risk factors of age, BMI, and gender had an area under the curve (AUC)/C - statistic of 85.6% while that of the African American PRS and five PCs was 58.6% (Figure 2). The AUC for prediction of type 2 diabetes increased by 0.6% with the addition of the PRS to the conventional risk factors.

**Figure 2.**
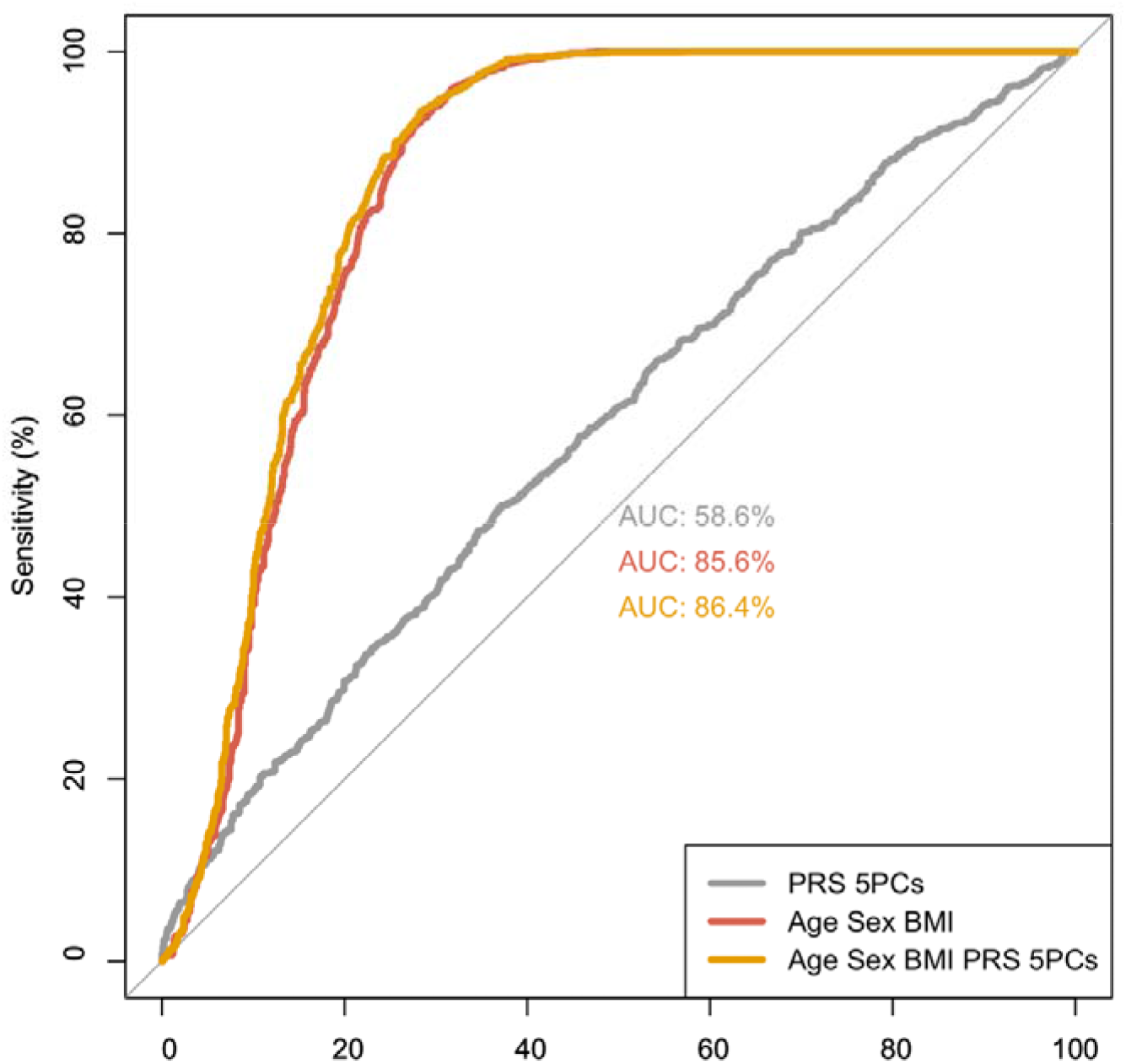
Receiver operating curves for the African Americans derived PRS and conventional risk factors for the prediction of type 2 diabetes in the South African Zulu study. Abbreviations; AUC= area under the curve, 5PCs = five principal components.

## Conclusions

Our study set out to assess the predictive value of type 2 diabetes PRS in continental Africans. In this study, we set out to compare the polygenic prediction of African American, European and multi-ethnic PRSs for type 2 diabetes in continental Africans. The PRS with the best prediction was derived from an African American restricted GWAS(7). Participants in the 10^th^ decile of this PRS had a more than 3-fold increased risk of developing type 2 diabetes and were diagnosed 2.6 years earlier on average as compared to those in the first decile.

Limited studies of candidate SNP PRS have been performed in continental Africans. Previously we reported a genetic risk score with weights from Europeans that was associated with OR = 1.21, 95%CI (1.02-1.43) for type 2 diabetes in black South Africans(19). This GRS had an AUC of 0.665 together with conventional risk factors for type 2 diabetes (19). However, this study was limited due to the small sample size (n = 356), the availability of only genotyped SNPs, and the use of weights that were derived from European-only studies. In our current study we have substantially expanded the sample size (n = 1,690), enhanced genome coverage by imputing to 1000 Genomes and local African Ancestry whole genomes(18), and used a multi-ethnic discovery dataset GWAS that included 1.4 million individuals which included people of African American ancestry. All these factors helped to enhance the predictive capability of PRS in our current study.

Nonetheless, polygenic predictions of European derived PRS in Europeans are still higher compared to that of the African Americans in continental Africans(7). Notably, participants in the top decile of a European derived PRS have recently been reported to have a greater than 5-fold risk for developing type 2 diabetes compared to those in the first decile in Europeans(7). Failure to reach predictions denoted in Europeans might be due to that in our study, the African American derived PRS are from an admixed population group that is not representative of the genetic diversity and linkage disequilibrium patterns of continental Africans(13; 20). In addition, vast improvements in sizes of the European cohorts that are now over a million individuals, is indicative of substantial power compared to African diabetes cohorts that are still below the 10 thousand mark (21). More investments are thus required to increase the representation of continental Africans in GWAS of type 2 diabetes.

The utility of polygenic risk scores is an issue of paramount importance for clinical translation(6). The African American PRS, though it was predictive for type 2 diabetes in continental Africans, only improved the AUC of conventional risk factors by 0.6% and when combined with PCs its AUC was 58.6% while that of the conventional risk factors was 85.6%. Similarly, in a Swedish type 2 diabetes study, the European derived PRS increased the AUC by 1 % compared to conventional risk factors (22). Basing on the AUC, the prediction of type 2 diabetes in our study by conventional risk factors (age, sex and BMI) was more clinically useful, as it was greater than the accepted threshold of 80%(6). However, the use of AUC as a measure to evaluate clinical utility of polygenic prediction is being debated, as it is regarded as less sensitive metric(23). There are ongoing efforts to develop better metrics (24), nonetheless, findings from this study that people with type 2 diabetes and a high PRS are typically diagnosed at an earlier age and have a 3 fold risk of developing diabetes are of clinical importance. They may be useful in prevention and treatment of diabetes.

Our study was limited by the limited number of GWAS of type 2 diabetes of continental Africans. Nonetheless, the African American derived PRS improved disease classification in this population. The clumping and thresholding approach used to compute the genome-wide PRS did not account for environmental factors such as diet, exercise that might confound the predictive accuracy of these measures. The strengths of our study include validation of the African American PRS in the AADM study and the fact that we used GWAS summary statistics of varied ethnicities from the same study which minimized bias due to genotyping and GWAS study designs.

In summary, an African American derived PRS seems to be the best predictor for type 2 diabetes in continental Africans, as compared to a European and multiethnic PRS. More studies are required to determine whether using continental African GWAS might further enhance these predictions and reach a similar accuracy as in Europeans. Although the PRS prediction of diabetes had low specificity and sensitivity, patient stratification by PRS may prove clinically useful.

## Contribution statement

TC and SF conceptulised the study, performed the main analyses and wrote the first draft. KE and AA performed the replication analysis. All the authors read and provided critical feedback on the paper.

## Acknowledgements

TC is an international training fellow supported by the Wellcome Trust grant (214205/Z/18/Z). SF is an international Intermediate fellow funded by the Wellcome Trust grant (220740/Z/20/Z) at the MRC/UVRI and LSHTM. S.F. received support from NIH grant U01MH115485 and the Makerere University-Uganda Virus Research Institute Centre of Excellence for Infection and Immunity Research and Training (MUII).MUII is supported through the DELTAS Africa Initiative (grant 107743). The DELTAS Africa Initiative is an independent funding scheme of the African Academy of Sciences (AAS), Alliance for Accelerating Excellence in Sciencein Africa (AESA), and supported by the New Partnership for Africa’s Develop-ment Planning and Coordinating Agency (NEPAD Agency) with funding fromthe Wellcome Trust (107743) and the U.K. government. DG was supported by the British Heart Foundation Centre of Research Excellence (RE/18/4/34215) at Imperial College, and a National Institute for Health Research Clinical Lectureship (CL-2020-16-001) at St. George’s, University of London. The DCC was funded by Servier South Africa, the South African Sugar Association and the Victor Daitz Foundation. The AADM study was supported in part by the Intramural Research Program of the National Institutes of Health in the Center for Research on Genomics and Global Health (CRGGH). The CRGGH is supported by the National Human Genome Research Institute (NHGRI), the National Institute of Diabetes and Digestive and Kidney Diseases (NIDDK), the Center for Information Technology, and the Office of the Director at the National Institutes of Health (1ZIAHG200362). Support for participant recruitment and initial genetic studies of the AADM study was provided by NIH grant No. 3T37TW00041-03S2 from the Office of Research on Minority Health.

## Conflicts of interest

DG is employed part-time by Novo Nordisk and has received consultancy fees from Abbott Laboratories.

---

**Supplementary Table 1.** The predictive performance of selected polygenic risk scores.

| **Ethnicity** | **P value Threshold** | **Nagelkerke R2** | **P value** | **Number of SNPs** |
| --- | --- | --- | --- | --- |
| European | $5 \times 10^{-8}$ | 0.003 | 0.002 | 6335 |
| European (Best PRS) | 0.406 | 0.007 | 2.57e-06 | 1441315 |
| Multiethnic | $5 \times 10^{-8}$ | 0.007 | 2.65e-06 | 7809 |
| Multiethnic (Best PRS) | $3 \times 10^{-4}$ | 0.008 | 1.26e-06 | 41815 |
| African America (Best PRS) | $5 \times 10^{-8}$ | 0.011 | 1.80e-08 | 65 |

## References

1. Saeedi P, Petersohn I, Salpea P, Malanda B, Karuranga S, Unwin N, Colagiuri S, Guariguata L, Motala AA, Ogurtsova K, Shaw JE, Bright D, Williams R. Global and regional diabetes prevalence estimates for 2019 and projections for 2030 and 2045: Results from the International Diabetes Federation Diabetes Atlas, 9th edition. Diabetes research and clinical practice 2019;157:107843

2. Ekoru K, Doumatey A, Bentley AR, Chen G, Zhou J, Shriner D, Fasanmade O, Okafor G, Eghan B, Agyenim-Boateng K, Adeleye J, Balogun W, Amoah A, Acheampong J, Johnson T, Oli J, Adebamowo C, Collins F, Dunston G, Adeyemo A, Rotimi C. Type 2 diabetes complications and comorbidity in Sub-Saharan Africans. EClinicalMedicine 2019;16:30–41

3. Langenberg C, Lotta LA. Genomic insights into the causes of type 2 diabetes. Lancet 2018;391:2463–2474

4. Hansen T. Type 2 diabetes mellitus--a multifactorial disease. Ann Univ Mariae Curie Sklodowska Med 2002;57:544–549

5. Diabetes Prevention Program Research G. Long-term effects of lifestyle intervention or metformin on diabetes development and microvascular complications over 15-year followup: the Diabetes Prevention Program Outcomes Study. The lancet Diabetes & endocrinology 2015;3:866–875

6. McCarthy MI, Mahajan A. The value of genetic risk scores in precision medicine for diabetes. Expert Review of Precision Medicine and Drug Development 2018;3:279–281

7. Vujkovic M, Keaton JM, Lynch JA, Miller DR, Zhou J, Tcheandjieu C, Huffman JE, Assimes TL, Lorenz K, Zhu X, Hilliard AT, Judy RL, Huang J, Lee KM, Klarin D, Pyarajan S, Danesh J, Melander O, Rasheed A, Mallick NH, Hameed S, Qureshi IH, Afzal MN, Malik U, Jalal A, Abbas S, Sheng X, Gao L, Kaestner KH, Susztak K, Sun YV, DuVall SL, Cho K, Lee JS, Gaziano JM, Phillips LS, Meigs JB, Reaven PD, Wilson PW, Edwards TL, Rader DJ, Damrauer SM, O’Donnell CJ, Tsao PS, Atkinson MA, Powers AC, Naji A, Kaestner KH, Abecasis GR, Baras A, Cantor MN, Coppola G, Economides AN, Lotta LA, Overton JD, Reid JG, Shuldiner AR, Beechert C, Forsythe C, Fuller ED, Gu Z, Lattari M, Lopez AE, Schleicher TD, Padilla MS, Toledo K, Widom L, Wolf SE, Pradhan M, Manoochehri K, Ulloa RH, Bai X, Balasubramanian S, Barnard L, Blumenfeld AL, Eom G, Habegger L, Hawes A, Khalid S, Maxwell EK, Salerno WJ, Staples JC, Yadav A, Jones MB, Mitnaul LJ, Aguayo SM, Ahuja SK, Ballas ZK, Bhushan S, Boyko EJ, Cohen DM, Concato J, Constans JI, Dellitalia LJ, Fayad JM, Fernando RS, Florez HJ, Gaddy MA, Gappy SS, Gibson G, Godschalk M, Greco JA, Gupta S, Gutierrez S, Hammer KD, Hamner MB, Harley JB, Hung AM, Huq M, Hurley RA, Iruvanti PR, Ivins DJ, Jacono FJ, Jhala DN, Kaminsky LS, Kinlay S, Klein JB, Liangpunsakul S, Lichy JH, Mastorides SM, Mathew RO, Mattocks KM, McArdle R, Meyer PN, Meyer U, Moorman JP, Morgan TR, Murdoch M, Nguyen X-MT, Okusaga OO, Oursler K-AK, Ratcliffe NR, Rauchman MI, Robey RB, Ross GW, Servatius RJ, Sharma SC, Sherman SE, Sonel E, Sriram P, Stapley T, Striker RT, Tandon N, Villareal G, Wallbom AS, Wells JM, Whittle JC, Whooley MA, Xu J, Yeh S-S, Aslan M, Brewer JV, Brophy MT, Connor T, Argyres DP, Do NV, Hauser ER, Humphries DE, Selva LE, Shayan S, Stephens B, Whitbourne SB, Zhao H, Moser J, Beckham JC, Breeling JL, Romero JPC, Huang GD, Ramoni RB, Pyarajan S, Sun YV, Cho K, Wilson PW, O’Donnell CJ, Tsao PS, Chang K-M, Gaziano JM, Muralidhar S, Chang K-M, Voight BF, Saleheen D, The HC, Regeneron Genetics C, Program VAMV. Discovery of 318 new risk loci for type 2 diabetes and related vascular outcomes among 1.4 million participants in a multi-ancestry meta-analysis. Nature genetics 2020;52:680–691

8. Martin AR, Kanai M, Kamatani Y, Okada Y, Neale BM, Daly MJ. Clinical use of current polygenic risk scores may exacerbate health disparities. Nature genetics 2019;51:584–591

9. Doumatey AP, Ekoru K, Adeyemo A, Rotimi CN. Genetic Basis of Obesity and Type 2 Diabetes in Africans: Impact on Precision Medicine. Curr Diabetes Rep 2019;19:105

10. Gaziano JM, Concato J, Brophy M, Fiore L, Pyarajan S, Breeling J, Whitbourne S, Deen J, Shannon C, Humphries D, Guarino P, Aslan M, Anderson D, LaFleur R, Hammond T, Schaa K, Moser J, Huang G, Muralidhar S, Przygodzki R, O’Leary TJ. Million Veteran Program: A mega-biobank to study genetic influences on health and disease. Journal of Clinical Epidemiology 2016;70:214–223

11. Chen J, Sun M, Adeyemo A, Pirie F, Carstensen T, Pomilla C, Doumatey AP, Chen G, Young EH, Sandhu M, Morris AP, Barroso I, McCarthy MI, Mahajan A, Wheeler E, Rotimi CN, Motala AA. Genome-wide association study of type 2 diabetes in Africa. Diabetologia 2019;62:1204–1211

12. Márquez-Luna C, Loh PR, Price AL. Multiethnic polygenic risk scores improve risk prediction in diverse populations. Genet Epidemiol 2017;41:811–823

13. Zakharia F, Basu A, Absher D, Assimes TL, Go AS, Hlatky MA, Iribarren C, Knowles JW, Li J, Narasimhan B, Sidney S, Southwick A, Myers RM, Quertermous T, Risch N, Tang H. Characterizing the admixed African ancestry of African Americans. Genome Biology 2009; 10: R141

14. Hird TR, Young EH, Pirie FJ, Riha J, Esterhuizen TM, O’Leary B, McCarthy MI, Sandhu MS, Motala AA. Study profile: the Durban Diabetes Study (DDS): a platform for chronic disease research. Global Health, Epidemiology and Genomics 2016;1:e2

15. Rotimi CN, Chen G, Adeyemo AA, Furbert-Harris P, Parish-Gause D, Zhou J, Berg K, Adegoke O, Amoah A, Owusu S, Acheampong J, Agyenim-Boateng K, Eghan BA, Jr., Oli J, Okafor G, Ofoegbu E, Osotimehin B, Abbiyesuku F, Johnson T, Rufus T, Fasanmade O, Kittles R, Daniel H, Chen Y, Dunston G, Collins FS, Africa America Diabetes Mellitus S. A genomewide search for type 2 diabetes susceptibility genes in West Africans: the Africa America Diabetes Mellitus (AADM) Study. Diabetes 2004;53:838–841

16. Rotimi CN, Dunston GM, Berg K, Akinsete O, Amoah A, Owusu S, Acheampong J, Boateng K, Oli J, Okafor G, Onyenekwe B, Osotimehin B, Abbiyesuku F, Johnson T, Fasanmade O, Furbert-Harris P, Kittles R, Vekich M, Adegoke O, Bonney G, Collins F. In Search of Susceptibility Genes for Type 2 Diabetes in West Africa: The Design and Results of the First Phase of the AADM Study. Annals of Epidemiology 2001;11:51–58

17. Adeyemo AA, Tekola-Ayele F, Doumatey AP, Bentley AR, Chen G, Huang H, Zhou J, Shriner D, Fasanmade O, Okafor G, Eghan B, Jr., Agyenim-Boateng K, Adeleye J, Balogun W, Elkahloun A, Chandrasekharappa S, Owusu S, Amoah A, Acheampong J, Johnson T, Oli J, Adebamowo C, Collins F, Dunston G, Rotimi CN. Evaluation of Genome Wide Association Study Associated Type 2 Diabetes Susceptibility Loci in Sub Saharan Africans. Front Genet 2015;6:335

18. Gurdasani D, Carstensen T, Fatumo S, Chen G, Franklin CS, Prado-Martinez J, Bouman H, Abascal F, Haber M, Tachmazidou I, Mathieson I, Ekoru K, DeGorter MK, Nsubuga RN, Finan C, Wheeler E, Chen L, Cooper DN, Schiffels S, Chen Y, Ritchie GRS, Pollard MO, Fortune MD, Mentzer AJ, Garrison E, Bergström A, Hatzikotoulas K, Adeyemo A, Doumatey A, Elding H, Wain LV, Ehret G, Auer PL, Kooperberg CL, Reiner AP, Franceschini N, Maher D, Montgomery SB, Kadie C, Widmer C, Xue Y, Seeley J, Asiki G, Kamali A, Young EH, Pomilla C, Soranzo N, Zeggini E, Pirie F, Morris AP, Heckerman D, Tyler-Smith C, Motala AA, Rotimi C, Kaleebu P, Barroso I, Sandhu MS. Uganda Genome Resource Enables Insights into Population History and Genomic Discovery in Africa. Cell 2019;179:984–1002.e1036

19. Chikowore T, van Zyl T, Feskens EJ, Conradie KR. Predictive utility of a genetic risk score of common variants associated with type 2 diabetes in a black South African population. Diabetes research and clinical practice 2016;122:1–8

20. Choudhury A, Aron S, Botigué LR, Sengupta D, Botha G, Bensellak T, Wells G, Kumuthini J, Shriner D, Fakim YJ, Ghoorah AW, Dareng E, Odia T, Falola O, Adebiyi E, Hazelhurst S, Mazandu G, Nyangiri OA, Mbiyavanga M, Benkahla A, Kassim SK, Mulder N, Adebamowo SN, Chimusa ER, Muzny D, Metcalf G, Gibbs RA, Matovu E, Bucheton B, Hertz-Fowler C, Koffi M, Macleod A, Mumba-Ngoyi D, Noyes H, Nyangiri OA, Simo G, Simuunza M, Rotimi C, Ramsay M, Choudhury A, Aron S, Botigué L, Sengupta D, Botha G, Bensellak T, Wells G, Kumuthini J, Shriner D, Fakim YJ, Ghoorah AW, Dareng E, Odia T, Falola O, Adebiyi E, Hazelhurst S, Mazandu G, Nyangiri OA, Mbiyavanga M, Benkahla A, Kassim SK, Mulder N, Adebamowo SN, Chimusa ER, Rotimi C, Ramsay M, Adeyemo AA, Lombard Z, Hanchard NA, Adebamowo C, Agongo G, Boua RP, Oduro A, Sorgho H, Landouré G, Cissé L, Diarra S, Samassékou O, Anabwani G, Matshaba M, Joloba M, Kekitiinwa A, Mardon G, Mpoloka SW, Kyobe S, Mlotshwa B, Mwesigwa S, Retshabile G, Williams L, Wonkam A, Moussa A, Adu D, Ojo A, Burke D, Salako BO, Matovu E, Bucheton B, Hertz-Fowler C, Koffi M, Macleod A, Mumba-Ngoyi D, Noyes H, Nyangiri OA, Simo G, Simuunza M, Awadalla P, Bruat V, Gbeha E, Adeyemo AA, Lombard Z, Hanchard NA, Trypano GENRG, Consortium HA. High-depth African genomes inform human migration and health. Nature 2020;586:741–748

21. Khera AV, Chaffin M, Aragam KG, Haas ME, Roselli C, Choi SH, Natarajan P, Lander ES, Lubitz SA, Ellinor PT, Kathiresan S. Genome-wide polygenic scores for common diseases identify individuals with risk equivalent to monogenic mutations. Nature genetics 2018;50:1219–1224

22. Lyssenko V, Laakso M. Genetic Screening for the Risk of Type 2 Diabetes. Worthless or valuable? 2013;36:S120–S126

23. Cook NR. Use and misuse of the receiver operating characteristic curve in risk prediction. Circulation 2007;115:928–935

24. Baker SG. Metrics for Evaluating Polygenic Risk Scores. JNCI Cancer Spectrum 2020;5

